# Context-Dependent Reactive Antipredator Behavior of Chacma Baboons *(Papio ursinus)* Amidst Predator Recovery

**DOI:** 10.64898/2026.04.05.716544

**Authors:** Sophia M. Van Cuylenborg, Nicholas S. Wright, Meredith S. Palmer, Susana Carvalho, Kaitlyn M. Gaynor

## Abstract

Predation is a driving force in the ecology and evolution of prey, and primates exhibit diverse anti-predator strategies for minimizing risk. Because these behaviors can be costly, individuals must balance costs and benefits when responding to perceived threats. The cognitive capacity and behavioral plasticity of baboons make them an ideal taxon for studying the context-dependent variation in anti-predator strategies. Here, we used an autonomous, motion-activated playback experiment to study the behavioral responses of chacma baboons (*Papio ursinus griseipes*) to simulated predator encounters in Gorongosa National Park, Mozambique. We compared responses in 2021, when predator densities were relatively low, to responses in 2024, after predation increased due to lion (*Panthera leo*) population recovery and African wild dog (*Lycaon pictus)* reintroduction. We compared flight and vigilance responses to vocalizations of these common predators with responses to leopard (*Panthera pardus*), historically a key predator; spotted hyena (*Crocuta crocuta*), a rare predator; and cheetah (*Acinonyx jubatus*), absent historically and currently. We also assessed how responses varied with habitat, age-sex class, presence of offspring, and group size. Across 916 predator playbacks, baboons fled in 19% and displayed vigilance in 71% of trials. When predator density was higher, baboons displayed weakened antipredator responses, consistent with the risk allocation hypothesis. Baboons were more likely to flee in response to lion and wild dog cues. Juveniles fled more frequently than other demographic classes, while adult females with offspring were more vigilant. Overall, responses were highly heterogeneous, reflecting the substantial intraspecific variation and behavioral flexibility characteristic of baboons.

## 1. Introduction

Predation is the leading cause of mortality in many primate species and therefore plays a key role in regulating their populations (Hill et al., 2019; Preisser et al., 2005) and shaping their behavior across ecological and evolutionary time scales (Shultz et al., 2011). Primates display a wide range of antipredator strategies, and their advanced cognitive capacity confers a high degree of behavioral plasticity, allowing them to fine-tune their responses in dynamic environments (Dall et al., 2005). These strategies can be divided into proactive behaviors that reduce the likelihood of encountering a predator and reactive behaviors utilized once predators are detected (Creel, 2018). Reactive behaviors, which are performed later in the predation sequence, have particularly high stakes given the imminent threat of death. The nature and intensity of reactive antipredator responses performed vary with many factors, including habitat structure, predator density and identity, and individual prey traits (Ouattara et al., 2009; Fornof et al., 2023).

Antipredator behaviors entail complex trade-offs between lowering predation risk and incurring energetic or opportunity costs, such as reduced foraging and mating. Flight is a common but energetically demanding reactive response, involving rapid movement away from a perceived threat (Ydenberg & Dill, 1986). Alternatively, prey can remain in place and engage in reactive vigilance, surveying their environment to assess relative danger (Lima, 1992). Vigilance is less energetically costly than flight, but is associated with an opportunity cost, given trade-offs with foraging (Brown, 1999). Although both behaviors lower the probability of predation, their frequent or prolonged use can be prohibitively costly if predator encounters are frequent (Ferrari et al., 2009). The “risk allocation hypothesis” therefore paradoxically suggests that as the background level of risk increases, prey allocate less time and energy towards antipredator behaviors (Lima & Bednekoff, 1999).

For prey facing risk from multiple predators, the strength of antipredator behavior towards a given species should vary by experience with that predator over both evolutionary and ecological time scales (Preisser et al., 2007; Sih et al., 1998). Given that predation imposes strong selective pressure on prey, prey should exhibit stronger perception of and response to risk from predators that historically accounted for a greater share of mortality. When predation risk relaxes on ecological time scales (e.g., following local predator extirpation), prey may nonetheless retain risk perception of their “ghosts of predators past” (Blumstein 2006). However, as maintaining unnecessary defenses is costly, antipredator behavior may weaken over time (Blumstein & Daniel, 2005; Jolly et al., 2018). If re-exposed to predators, naïve prey can rapidly learn to associate novel stimuli with risk, especially when the cues are paired with predation attempts or alarm calls of conspecifics or heterospecifics (Griffin et al., 2000). In social primates, individual learned responses can be transmitted within social groups or populations (Griffin & Evans, 2003). Consequently, the expression of anti-predator behavior reflects the interaction between evolutionary history and individual and social learning, producing variable responses across dynamic predation landscapes.

Demographic, social, and environmental characteristics also strongly mediate antipredator responses in primates. Juveniles are typically the most vulnerable age class because of their small size, inexperience, and limited locomotor abilities (Hill et al., 2019). As a result, juveniles often show more intense antipredator responses or rely heavily on parental protection (Mohr et al., 2023). In sexually dimorphic species, sex differences in body size and physical condition further influence vulnerability to predation. Larger males are generally more conspicuous but better able to deter or escape predators, while smaller or reproductively encumbered females may face greater risk due to reduced mobility or energy reserves (Caro, 2005; Pettorelli et al., 2011). Additionally, group size can influence individual risk assessment: primates often increase vigilance when alone or in smaller groups, whereas in larger groups, the dilution and “many-eyes” effects (shared vigilance) can reduce individual vigilance (Dehn, 1990; Lehtonen & Jaatinen, 2016; Pulliam, 1973). Finally, habitat structure influences how primates respond to predators: the utility of vigilance may be greater in open habitats, where detection is easier, but risk may be higher in closed habitats where visibility is limited, thus necessitating greater vigilance (Lima & Dill, 1990; Fornof et al., 2023). However, primates are often more adapted to arboreal environments than their predators (Ouattara et al., 2009), and trees may therefore provide a spatial refuge from terrestrial predators that are not available in open habitats.

Baboons offer a useful model for studying primate antipredator behavior. Baboons are highly terrestrial and are therefore vulnerable to a large suite of mammalian carnivores, including wild dogs (*Lycaon pictus*), lions (*Panthera leo*), and spotted hyenas (*Crocuta crocuta*; Galán-Acedo et al., 2019; Dzingwena et al., 2025; Hammond et al., 2025; van der Meer et al., 2019). Their primary predators, however, are leopards (*Panthera pardus*), which are unique among mammalian carnivores in their ability to hunt baboons both terrestrially and arboreally (Cowlishaw, 1997), particularly at night, when baboons sleep in the trees (Cowlishaw, 1994; Isbell et al., 2018). Within baboon groups, predation risk varies markedly among individuals: as highly social animals with a dominance structure that has defined spatial roles for males, females, and juveniles, males often occupy riskier periphereral positions (Lehmann & Ross, 2011; van Schaik et al., 2022). Further, baboons exhibit significant behavioral plasticity (Fehlmann et al., 2017) and cognitive capacity (Fagot & Cook, 2006), traits that may aid them in responding quickly to shifts in predation risk via individual or group learning mechanisms (Hammond et al., 2022).

Here, we investigated how predation risk, predator characteristics, and social and environmental context shape the flight and vigilance behavior of chacma baboons (*Papio ursinus griseipes*) in Gorongosa National Park, Mozambique. We conducted a playback experiment using Automated Behavioral Response (ABR) systems, which employ a camera trap to record baboon behavioral responses to automated audio playbacks of predator vocalizations and control sounds (Palmer et al. 2022; Smith et al., 2020; Suraci et al., 2017). ABRs have been used to study fear responses of ungulates to apex predators and megaherbivores (Fletcher et al., 2023; Rigoudy et al., 2022) and the impacts of humans on wildlife (Zeller et al., 2024). Further, they can address some of the major challenges in studying primate antipredator behavior, including the rarity of observing primate predation events (Lin et al., 2020) and the influence of human observers on the behavior of both primates and their predators (Nowak et al., 2014; Allan et al., 2020; Iredale et al., 2010). ABRs offer a promising answer to the call for an increased use of indirect methods to studying primate-predator interactions (LaBarge et al. 2020).

Gorongosa National Park provides a unique natural experiment for understanding the effects of predation on baboon behavior. Gorongosa’s large carnivores historically included lions, African wild dogs, leopards, and spotted hyenas. During Mozambique’s civil conflict (1977–1992), all large carnivores except for lions were extirpated (Pringle & Gonçalves, 2022), and the population of lions was greatly reduced (Bouley et al., 2018). The park has since reintroduced all large carnivore species, and during the period of our study (2021 to 2024), populations of lions and wild dogs rose, and leopards and spotted hyena were first reintroduced (Table S1). We evaluated responses of baboons to cues from each of Gorongosa’s large carnivore species, as well as cheetah (*Acinonyx jubatus*), a novel species that was never present in Gorongosa.

We hypothesize that baboon reactive antipredator behavior is influenced by several factors: a) predator exposure (year as a proxy for relative predator density), b) predator identity, c) prey demographics, d) group size, and e) habitat type. Under the learned responses hypothesis, baboons should exhibit stronger antipredator responses as exposure to predators increases between 2021 and 2024. Alternatively, the risk allocation hypothesis predicts that baboons exhibit weaker antipredator responses in 2024, as exposure to predators increases. We predicted that baboons would respond most strongly to cues from leopards, their primary predator over evolutionary history, given greater perceived risk, whereas they would respond less strongly to cues from cheetahs, as they are not a historically important predator of baboons and are absent from the study area. Given that vulnerability should shape antipredator responses, we predicted that adult female baboons with offspring would exhibit stronger antipredator responses than other demographics, and that responses should be weaker when individuals are in larger groups. Finally, given that closed habitats offer more arboreal refugia from predators but also obscure visibility, we predicted stronger flight and vigilance in closed habitats relative to open habitats. While we were also interested in understanding interactions among these mediators of behavior (e.g., how the effects of habitat and year varied with predator identity), sample size constraints precluded these analyses, so we instead focused on evaluating the independent effects of ecological and social factors on reactive baboon antipredator responses.

## 2. Methods

### 2.1 Ethical Note

This study was approved by The University of British Columbia Animal Care Committee (A24-0038) and by Princeton University’s Institutional Animal Care and Use Committee (IACUC #2119F-18). The research complied with the legal requirements of Mozambique, and a research permit was granted by Gorongosa National Park’s Department of Scientific Services. All procedures adhered to the American Society of Primatologists’ Principles for the Ethical Treatment of Non-Human Primates.

### 2.2 Study System

Gorongosa National Park is a 4,000 km² protected area in central Mozambique, established in 1960 (Figure 1). The park’s wildlife population was severely depressed during Mozambique’s civil conflict, amidst rampant bushmeat hunting, with large carnivore populations particularly affected by both snaring and prey loss (Daskin & Pringle, 2018; Stalmans et al., 2019). Following the end of the conflict, and facing relaxed predation pressure, many non-carnivore large mammal species recovered quickly (Stalmans et al., 2019). Currently, chacma baboons *(Papio ursinus)* are among the most abundant mammal species in the park (Gaynor et al., 2021; Martinez et al., 2019; Carvalho et al., 2025), and there are now over 200 troops (Stalmans and Peel, 2024). In recent years, the park has since reintroduced the extirpated carnivore species (Table S1), beginning with the translocation of 45 African wild dogs starting in 2018 (Bouley et al., 2021; Pringle & Gonçalves, 2022). African wild dogs had a minimum estimated population size of 130 in 2021, which increased to over 250 in 2024 (Table S1). In 2021, four leopards were introduced to the park (numbering 7 in 2024). In 2022, spotted hyena reintroduction began, with an estimated 23 individuals in the park as of 2024 (Table S1). Lion populations, which were extremely depressed post-conflict (Bouley et al., 2018), were estimated to have a minimum population size of 130 in 2021, which increased to 245 in 2024 (Table S1).

**Figure 1.**
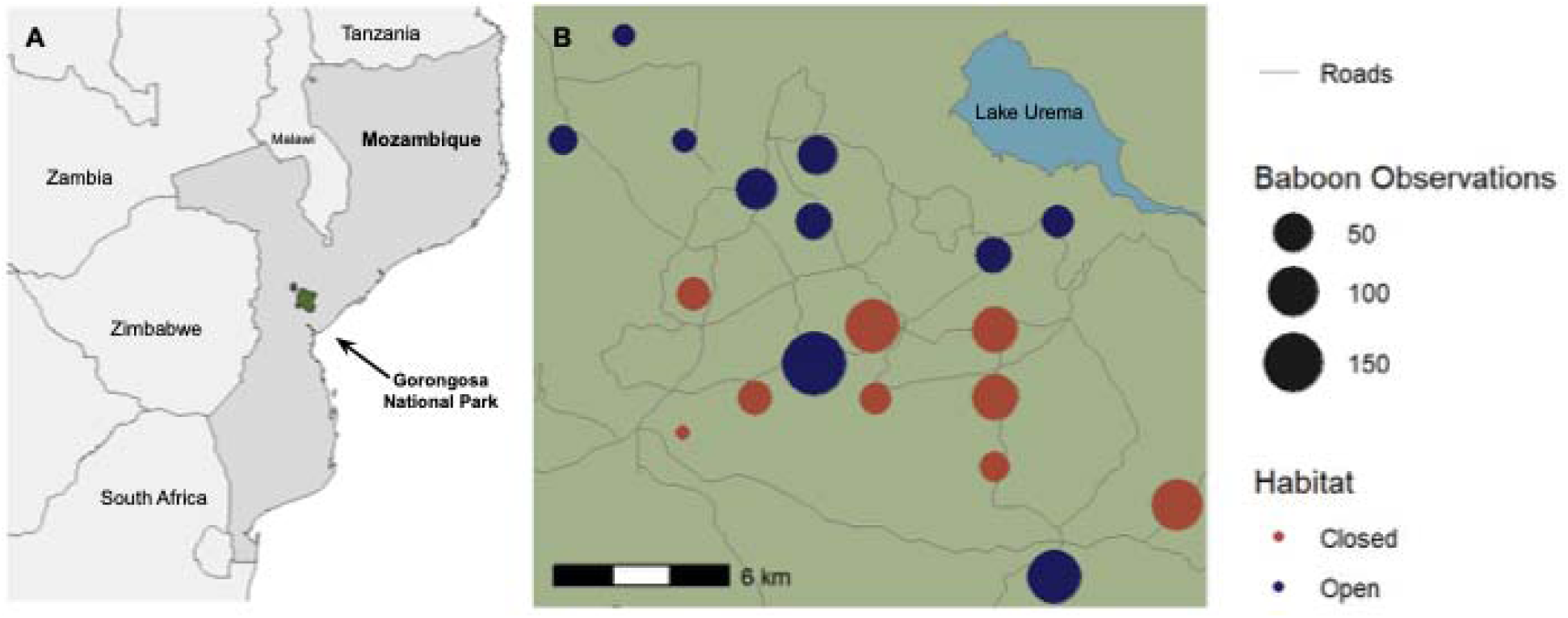
Location of Gorongosa National Park in Mozambique (left) and location of Automated Behavioral Response (ABR) system deployments within the park (right). Each circle represents the location of an ABR site, colored by habitat. The size of the circle represents the number of baboon videos captured at the site in both 2021 and 2024.

### 2.3 Study Design

We set up the Automated Behavioral Response systems (ABRs) following the methodology in Palmer et al. (2022). Each system incorporated a Browning Strike Force Pro camera trap and a Freaklabs BoomBox (audio system; https://boomboxproject.org/). We programmed the soundboard with predator vocalizations from five mammalian species (leopard, lion, spotted hyena, cheetah, and African wild dog) and control sounds from two avian species (the southern ground hornbill, *Bucorvus leadbeateri*, and the African hoopoe, *Upupa africana*). For each predator species, we included a variety of representative vocalizations, such as growls and snarls, to capture acoustic variability. The chosen species represent the largest mammalian carnivores present in the park at the time of the study, as well as cheetahs, which represent a non-native species never present in the study system. The avian control sounds allowed us to evaluate how much of the behavioral response was driven by the sudden loud sound, versus the predator cue specifically.

We edited all sounds for consistent amplitude and quality using Audacity (v. 2.1.0; Audacity Team, 2014). Following Suraci et al. (2017), we standardized temporal properties by visually inspecting spectrograms and waveforms, and we used t-tests to confirm that frequency characteristics (peak, minimum, maximum, range) did not differ significantly between predator and control groups. We adjusted the playbacks to a consistent peak sound pressure level of 80 dB at 1 m, measured with a Radioshack 33-2055 Digital Sound Level Meter set to fast response and C-weighting. Each video lasted 30 seconds, followed by a 30-second rest period before another trigger could re-activate the ABR system. Each device played a randomized playlist of all auditory cues, and the order of the cues varied across cameras.

### 2.3 Data Collection

We deployed ABRs at 19 sites in the core road network of Gorongosa National Park, with 10 in open habitats (floodplains, grasslands, or open savanna) and 9 in closed habitats (woodlands or dense shrublands) (Figure 1). The ABRs were each ∼100m from established camera trap sites that were part of a long-term grid, and the sites were chosen based on prior animal detection rates to ensure adequate sample sizes (Gaynor et al. 2021). We deployed the ABRs in the same locations in two dry seasons: June–August 2021, when predator density was still low, and July–August 2024, when predator density had increased. Two of the sites were excluded in 2024 due to inaccessibility. To minimize habituation to audio cues from repeated exposure, each sampling period lasted 2–3 months.

There were a total of 1,462 ABR videos of chacma baboons (n=967 from 2021 and n=495 from 2024). Two ABR sites had far greater baboon detections (one open habitat site had 485 detections in 2021, and 120 in 2024; one closed habitat site had 109 detections in 2024) compared to the other sites (mean =19 detections in a year). The large number of detections made manual annotation of all videos infeasible, so for these three site-years we selected a random subset of 88 videos, as this was the number of videos for the site-year with the third-highest detections. We also excluded 96 videos in which the auditory cue did not play properly. In total, we analyzed a total of 916 ABR videos (n=517 in 2021; n=399 in 2024).

After developing an ethogram (Table S2), we classified behaviors frame-by-frame using the Computer Vision Analysis Tool (CVAT; Cvat.ai Corporation 2025). To minimize bias, we annotated videos without audio so that the observer did not know whether the video contained a control or predator sound. For videos featuring multiple individuals, we identified the focal subject as the individual closest to the camera, at the center of the frame - presumably the one who triggered the camera. For analysis, we merged the behaviors into four (4) classes to simplify inference: vigilant, non-vigilant, occluded, and flight behavior (Table S2; Figure 2). We reviewed each video a second time with sound to record the species of predator or control audio cue, group size (number of unique individuals appearing in the 30-second video), and prey demographic characteristics, including age (adult or juvenile), sex (male or female, for adults only), and presence of offspring (young juveniles that were carried by their mother).

**Figure 2.**
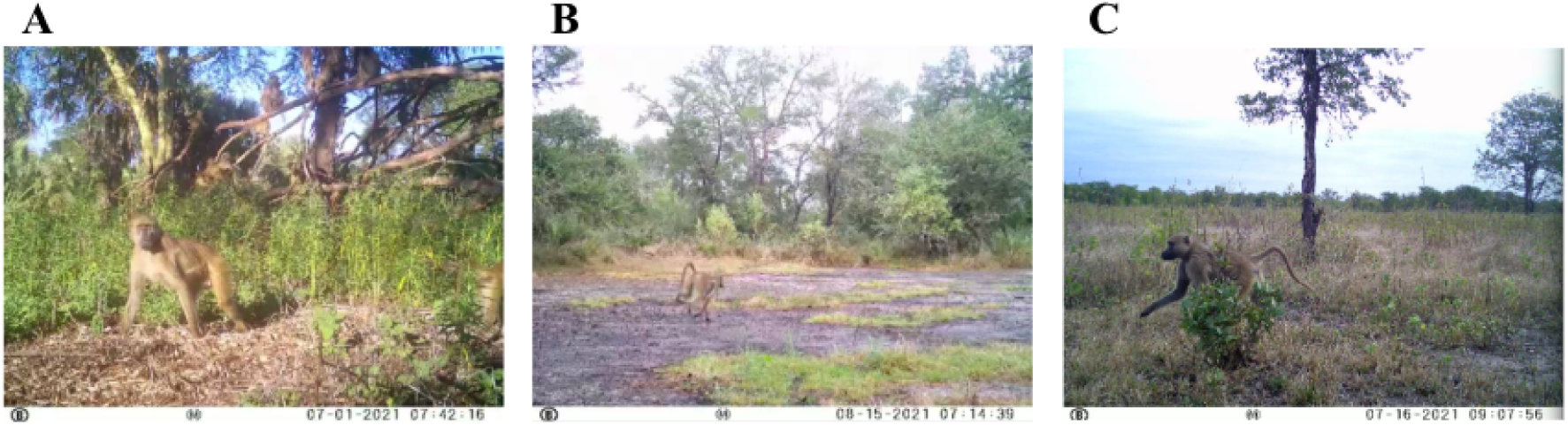
Chacma baboon behavior exemplars in response to simulated predator audio cues from our ABR study in Gorongosa National Park. (A) vigilance (B) flight (C) non-vigilant walking.

### 2.4 Data Analysis

Our two response variables of interest were (1) flight, a binary variable representing whether or not the baboon fled following the predator cue playback (at any point in the 30-second video), and (2) vigilance, the proportion of non-occluded frames in which the focal baboon was vigilant. We modeled flight using generalized linear mixed models (GLMM) with a binomial distribution. For vigilance, we used GLMMs with a beta distribution after transforming the proportion of vigilance using the Smithson & Verkuilen transformation (_____________) to accommodate values of 0 and 1 (Smithson & Verkuilen, 2006). For the vigilance analysis, we excluded videos in which baboons fled immediately (latency to flee = 0 seconds).

To confirm that baboons were responding to information in the predator cue rather than the sudden auditory stimulus itself, we first ran a single model for each response variable with a binary covariate representing whether the auditory cue was a control (avian vocalization) or treatment (predator vocalization) and included the ABR site as a random effect.

Then, to test our hypotheses about the factors mediating variation in flight and vigilance, we ran models using only the predator cue videos. Fixed effects included predator species (cheetah, hyena, leopard, lion, or wild dog), year (2020 or 2024), habitat (open or closed), group size, and day of study. We included day of study (number of days since the ABR was deployed in a given year) to account for potential effects of habituation to the cues over the course of the study. We also included the ABR site as a random effect in all models to account for site-level variation and potential pseudoreplication arising from repeated sampling of the same individual or troop. We performed all analyses using R version 4.4.3.

We used the ‘dredgè function from the MuMIn package (Bartoń, 2024) to generate all possible subsets of the global model, and ranked models based on the corrected Akaike Information Criterion (AICc). AICc balances model fit (via log-likelihood) with model complexity (number of parameters), and is more appropriate than AIC for small-to-moderate sample sizes (Burnham & Anderson, 2004). We used the ‘model.avg’ function in MuMIn to calculate an AIC-weighted parameter estimate, averaged across all models in which that parameter appeared. This approach accounts for uncertainty in model structure by allowing inference on multiple explanatory variables across a suite of plausible models, rather than relying on a single “best” or potentially overfit model.

We assessed the relative importance of covariates based on the sum of AIC weights across all models that included that variable and evaluated effect sizes by calculating a model-averaged regression coefficient and its 95% confidence interval (CI). This approach is appropriate for our data because we specified predictors a priori based on our hypotheses, and inference based on P-values would be inappropriate given model averaging. To determine the predicted effects of year, predator characteristics, prey demographics, and habitat on vigilance and flight probability, we used the ‘predict()’ function on the averaged model excluding random effects, holding other variables constant while varying one predictor of interest.

To explore the cumulative probability of flight as a function of time since cue playback, we used a Kaplan-Meier survival analysis and constructed survival curves. This approach estimates latency to flee while also accounting for censored observations (in which there was no flight observed in the video timeframe).

## 3 Results

Baboons fled from the predator auditory cues in 19% of trials (in contrast to 11% for the control cues), with a mean latency to flee of 1.08 seconds (standard deviation = 0.81 seconds) after the cue began to play. Baboons were significantly more likely to flee in response to predator cues than to the avian control cues (beta = 0.676, SE = 0.313, P = 0.031).

Of the baboons that did not immediately flee from the predator cues, 82% of them exhibited some amount of vigilance (vigilance was displayed in 71% of the predator cue trials overall). In response to the predator playbacks, baboons were vigilant for a mean of 43% of the time that they were in frame (standard deviation = 33%). In contrast, baboons were vigilant for a mean of 38% of the time in response to control cues (standard deviation = 34%). Baboons were significantly more vigilant in response to predator cues relative to the control (beta = 0.289, SE = 0.136, P = 0.034). Of all videos in which baboons exhibited vigilance, 16% of the videos included scanning, 39% staring, and 74% moving with vigilance (as described in the ethogram).

### 3.1 Flight predictors

The probability of flight in response to predator cues varied with environmental and social context (Figure 3A, Figure S1, Table S3). Age-sex class was the most important predictor of flight in baboons, with juveniles the most likely demographic to flee. Adult males and females with offspring were also slightly more likely to flee than adult females without offspring. Year was also an important predictor of flight: baboons were less likely to flee in 2024 than in 2021. There was also strong support for predator species as a predictor of flight, with the greatest flight responses elicited by lions, followed by wild dogs, leopards, hyenas, and cheetah. There was moderate support for group size, with larger group size associated with lower flight probability, although the effect was weak. There was also moderate support for habitat, with some evidence that baboons are more likely to flee in closed relative to open habitats. The day of study had the weakest support as a predictor of flight. Overall, our model explained a low proportion of the variation in flight behavior, with a conditional R² value of 0.162.

**Figure 3.**
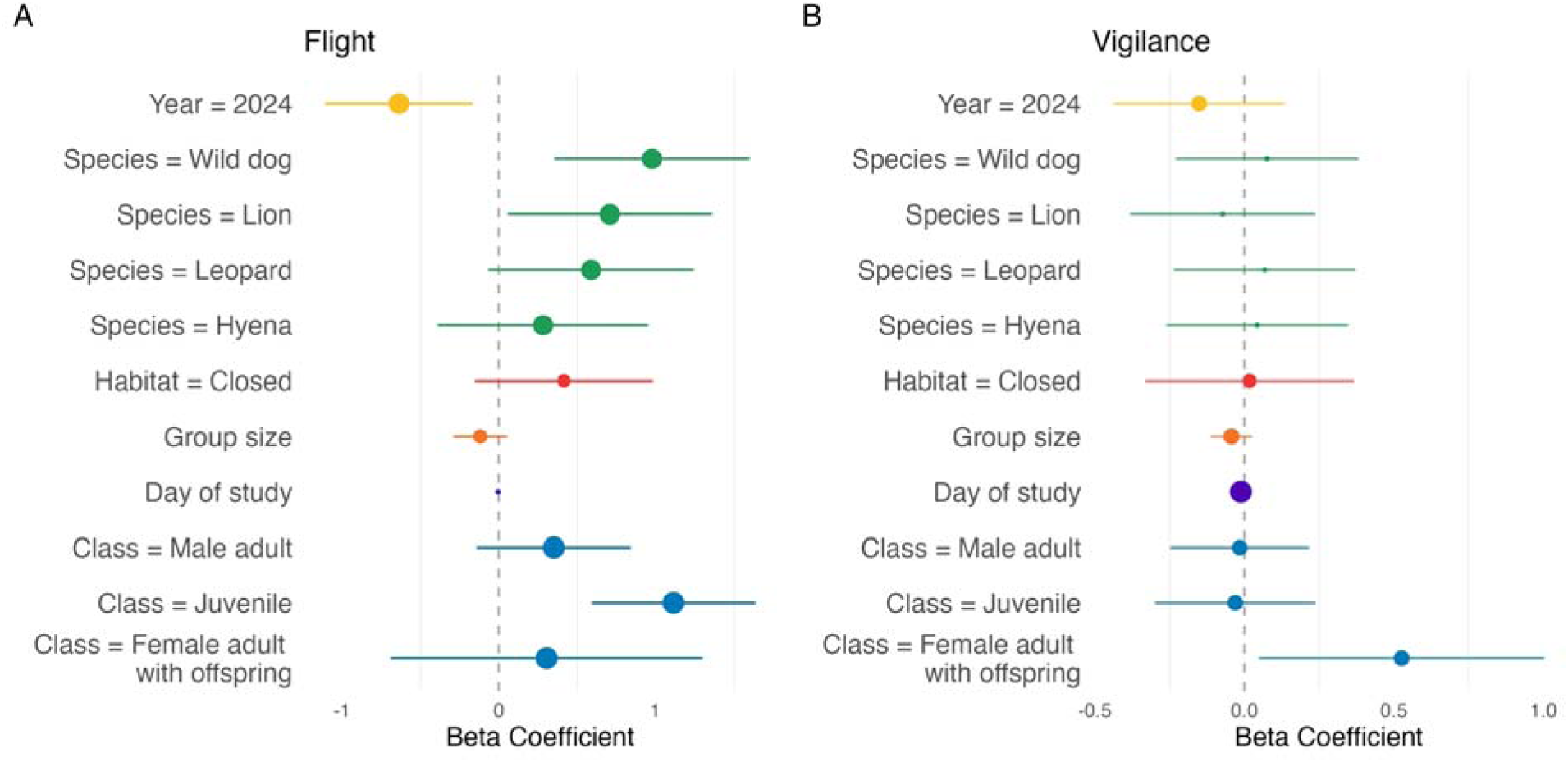
Effects of the environmental, social, and demographic context on A) flight and B) vigilance behavior of baboons in response to simulated auditory predator cues in Gorongosa National Park, Mozambique. The size of the point corresponds to the relative variable importance (sum of AIC weight for all models in which the covariate appears). Each covariate is represented by a unique color. Error bars correspond to 95% CIs. For the categorical variables, the reference categories are Year = 2021, Species = Cheetah, Habitat = Open, Class = Adult female (without offspring). Positive beta coefficients correspond to a positive effect of that covariate (relative to the reference category, for categorical variables) on A) probability of flight or B) percent time vigilant, while negative coefficients correspond to a negative effect.

Kaplan–Meier survival analysis indicated that flight is an immediate response when it occurs. Of the baboons that fled, 90% did so within 2 seconds of cue initiation. The subsequent flattening of the curve suggests that the probability of flight declined as the elapsed time from the cue increased (Figure 4).

**Figure 4.**
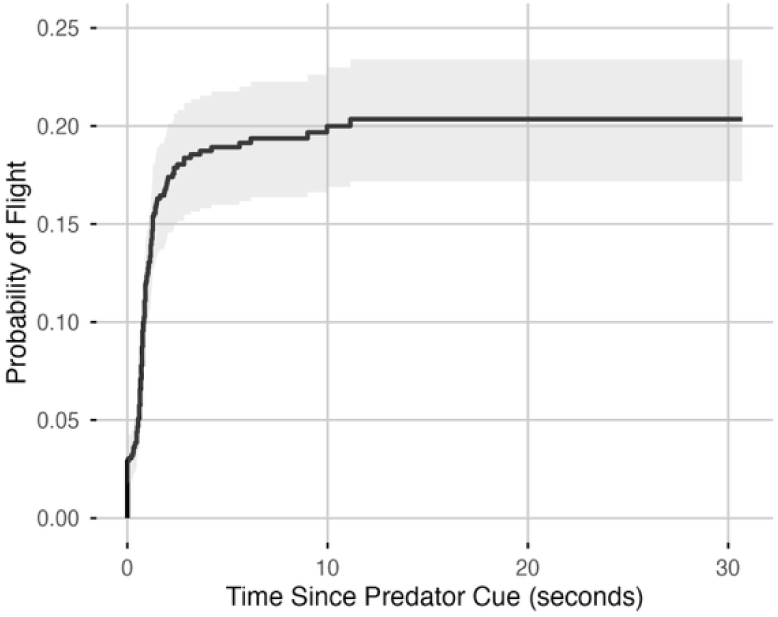
Kaplan–Meier survival curve showing the probability that baboons in Gorongosa National Park, Mozambique, fled with elapsed time since experimental predator audio cues. Shaded areas indicate 95% CIs.

### 3.2 Vigilance predictors

Few predictors of vigilance were strongly supported, based on overall AIC weights across models (Figure 3B, Figure S2, Table S4). The day of study was the most important predictor of vigilance, but its effect size was small, suggesting that vigilance decreased slightly over the study period each year. There was some evidence that baboons were less vigilant in 2024 than in 2021, with moderate support for year as a predictor. Age-sex class was also a moderately important predictor of vigilance, with female adults with offspring showing greater vigilance relative to the other classes. Group size was also supported, but as with flight, its negative effect on vigilance was small. Habitat had weak support and near-zero effect on vigilance. There was very little support for predator species as a predictor of vigilance, with all five species eliciting similar vigilance responses. Overall, our model explained a low proportion of the variation in vigilance, with a conditional R² value of 0.201.

## 4. Discussion

In response to simulated predator vocalizations, baboons in Gorongosa National Park exhibited heterogeneous anti-predator responses that varied by predator identity, year, and environmental and social context. Overall, baboons were far more likely to exhibit vigilance than flight: 71% of baboons were vigilant after the onset of the cue, while 19% fled. Vigilance enables ongoing assessment of a threat under uncertainty, and such information-gathering may be especially necessary when predator cues are novel, ambiguous, or rare, or in this case, when an auditory cue is decoupled from any visual or olfactory cues (Jones et al. 2024). In contrast, flight is a decisive response to perceived immediacy or certainty of danger, and when baboons fled, they typically did so immediately. The decision to flee was more strongly shaped by the environmental context and individual identity than vigilance, likely because the flexibility and lower cost of vigilance enable it to be more widely used across contexts.

Predator-dependent flight responses suggest that predators are not a functionally interchangeable trophic level, but instill varying levels of perceived risk to which prey respond accordingly. This idea of a “hierarchy of fear” echoes findings of a recent ungulate ABR study (Rigoudy et al. 2022). Baboons were more likely to flee in response to cues from wild dogs, which comprise the fastest-growing predator population in Gorongosa (Bouley et al., 2021; Table S1), and to lions, the other common predator in the park and the only large carnivore that was never locally extirpated (Bouley et al., 2018). The heightened response to wild dogs is likely because they are a relatively novel and common predator, given that they do not normally prey on baboons. Baboons also fled more from cues of leopards, the primary predators of baboons across savanna ecosystems (Cowlishaw, 1994). Given the low leopard density during the study, this antipredator behavior likely represents a hardwired evolutionary response. Baboons responded most weakly to cheetah cues, likely because this population of baboons lacks both recent and long-term experience with cheetahs as a predator (Anderson et al., 2006; Tinley, 1977). Thus, both experience and evolutionary history appeared to shape variation in behavioral responses to different predator species.

Baboons fled less frequently and were less vigilant in 2024 than in 2021, despite increases in predator densities in the intervening years. This weakening of anti-predator behaviors with greater predation risk is consistent with the risk allocation hypothesis, and is consistent with findings from other systems (Creel et al. 2008; Ferrari et al., 2009). Reduced anti-predator behavior in 2024 may reflect a shift towards strategic risk tolerance, where baboons weigh the relative value of immediate safety against the long-term needs of energy acquisition and rest (Lima, 1998). These findings echo previous studies on the ability of baboons to adjust to the dynamic and rapidly changing environment of Gorongosa, including evidence of plasticity in spatial behavior in response to both seasonal flooding (Lewis-Bevan et al. 2025) and extreme weather events (Beardmore-Heard et al. 2025).

Individual demographic characteristics influenced the strength of baboon antipredator behaviors. The least likely demographic to flee was adult females, which have the most experience with assessing predation risk in the troop’s home range, and may be less likely to equate auditory cues with risk in the absence of other cues (e.g., visual and olfactory cues, alarm calls). Juveniles were most likely to flee, consistent with their disproportionately high predation risk given their size, and lack of experience recognizing and assessing risk (Pettorelli et al., 2011). Adult females with offspring, which face a higher risk and cost of predation, were marginally more vigilant, consistent with patterns observed in other primates (Liu et al., 2024).

We found weak evidence that both flight and vigilance decreased when individuals were in larger groups. As group size increases, baboons may be less likely to flee and more likely to collectively mob a predator, given that mobbing is an effective defense strategy (Crofoot 2013). Although the ‘many eyes’ effect is common among prey species (Beauchamp et al., 2021), its evidence in primates is mixed, as vigilance is also heavily shaped by social monitoring and group dynamics (Gaynor & Cords, 2012; Allan et al., 2024). Notably, we were limited by our ability to count only the number of individuals that were in frame, which may not accurately reflect the actual group size, as chacma baboon troops can range from <10 to >150 individuals (Hammond et al. 2025). We also found weak evidence that baboons were more likely to flee in closed habitats than in open habitats as predicted, and our observations suggest that baboons often take advantage of refuge in trees to flee vertically. However, there was no effect of habitat type on vigilance, and trade-offs between visibility and concealment may minimize the effect of habitat on vigilance (Camp et al., 2013).

Our findings demonstrate a high degree of intraspecific variation, in which different individuals respond very differently to the same threat in the same environmental context (Cooper & Blumstein, 2015). Behavioral responses are complex and difficult to predict (Crane et al., 2024), and much of the variation in baboon reactive anti-predator behavior remains unexplained by the variables we examined. While we were unable to identify individual animals in our study, consistent individual responses to risk may be shaped by personality or behavioral syndromes, including boldness (Sih et al., 2004). In cognitively advanced and social species like baboons, individual variation may be further shaped by prior experience and social dynamics, and the interplay between personality and sociality. In wild chacma baboons, for example, boldness and anxiety predict propensity for social learning (Carter et al., 2014). Further, because baboons are a hierarchical species, dominance rank can influence access to resources and group dynamics, which in turn may affect individual responses to predators (Gesquiere et al. 2025; Fele et al. 2026). Future research should assess the role of individual variation alongside environmental context in shaping anti-predator behavior.

Our study used recordings of predator vocalizations to simulate a predation event, which, while effective in eliciting behavioral responses from prey, is not a perfect proxy for an actual encounter. Habituation is a potential concern with studies involving simulated predator cues, such as playback experiments, given that the cues are not reinforced by actual predator encounters. We found that vigilance decreased slightly over the course of the study, although the overall effect of habituation was minimal, and there was no effect of day on flight probability. However, this finding highlights that habituation may weaken effect sizes in experimental studies of anti-predator behavior in response to artificial predator cues. Furthermore, acoustic cues alone do not capture the full range of sensory information prey use to assess risk, including visual or olfactory signals (Munoz & Blumstein, 2012). Additionally, ambush predators like lions and leopards rely on stealth, and thus rarely vocalize when hunting. However, carnivore vocalizations for territorial advertisement, social communication, or intimidation still signal the presence of a nearby predator (Hettena et al. 2014). Despite these limitations, audio cues offer a controlled, repeatable, and non-invasive method for simulating predation risk (Smith et al., 2020). Future studies could incorporate multimodal cues to better capture the range of sensory information prey use to assess predation risk (Zuberbühler, 2007; Jordan & Ryan 2015).

This work contributes to an understanding of how baboons, as socially and cognitively complex prey species, balance competing ecological pressures and adapt to changing predator landscapes. Anti-predator behavior is modulated by the factors explored in this study–learned responses, predator identity, prey demographics, and habitat–and other unexplored factors (Ouattara et al. 2009). Further research is needed to fully understand and untangle these complex drivers of anti-predator behavior. Studying these behaviors provides critical insights into the role predators play in shaping the evolution of cognition, sociality, and life history strategies in primates.

## 5. Acknowledgments

This research was made possible by the permission of the Mozambican government and the support of Gorongosa National Park, particularly P. Muagura, M. Stalmans, and M. Lajas. We are grateful to the many rangers who offered their protection and expertise in the field. C.A. Wang and J. Plucinski at Freaklabs built the BoomBox Automated Behavioral Response system and helped us troubleshoot technical difficulties. S. Sun helped assemble and program the ABRs. M. Angela provided data on carnivore populations in the park. Funding for this work came from NSERC Discovery Grant (RPGIN-2022-03096) to KMG, a Sloan Research Fellowship (FG-2024-21781) to KMG, an NSF Postdoctoral Research Fellowship in Biology (Award No. 1810586) to MSP, and an Animal Behavior Society Student Research Grant to NSW. This work is part of the AI and Biodiversity Change (ABC) Global Climate Center, which is supported by the US National Science Foundation award No. 2330423 and Natural Sciences and Engineering Research Council of Canada award No. 585136. We thank the Gaynor Lab (UBC) and Pringle Lab (Princeton) for their feedback and support, particularly R. Pringle. We would like to acknowledge that The University of British Columbia, where much of this work took place, is located on the unceded traditional territories of the xʷməθkʷəy̓əm (Musqueam), Sḵwx̱wú7mesh (Squamish), and səlilwətaɬ (Tsleil-Waututh) Nations.

## Abbreviations

ABR system: Automated Behavioral Response system

