## Supplemental Information for "Context-Dependent Reactive Antipredator Behavior of Chacma Baboons *(Papio ursinus)* Amidst Predator Recovery"

### 7. Supporting Information

**Table S1. Annual large carnivore population estimates for Gorongosa National Park.** Abundance estimates represent minimum known population sizes based on individual monitoring by the park's veterinarian team.

| Species | 2017 | 2018 | 2019 | 2020 | 2021 | 2022 | 2023 | 2024 |
|---|---|---|---|---|---|---|---|---|
| African wild dog | 0 | 14 | 54 | 81 | 130 | 171 | 197 | 253 |
| Lion | Unknown* | Unknown* | 156 | 164 | 188 | 200 | 217 | 245 |
| Leopard | 0 | 1 | 1 | 2 | 6** | 5 | 5 | 7 |
| Spotted hyena | 0 | 0 | 0 | 0 | 0 | 6 | 14 | 23 |

*The lion population was estimated to be 104 in 2016, but no counts were conducted in 2017/18.

**During the 2021 sampling period for this study, there were four leopards in Gorongosa. Two more were reintroduced later that year.

**Figure S1. Relative flight frequency for baboons in response to simulated predator auditory cues in Gorongosa National Park, Mozambique, across A) predator cues, B) year, C) habitat, and D) baboon age and sex class.** Panels B-D exclude data from playbacks of control (avian) cues. Error bars correspond to 95% confidence intervals.

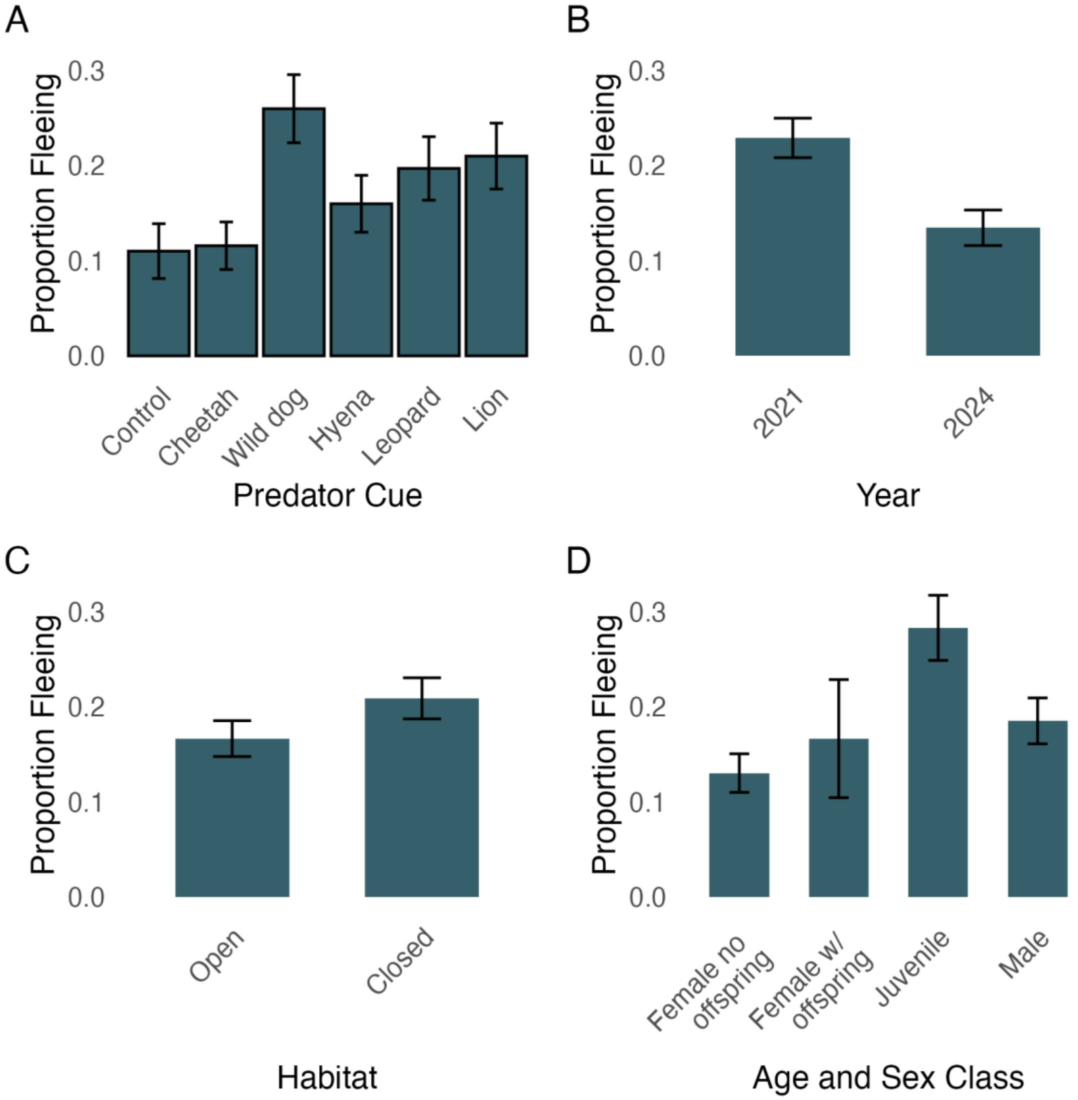

**Figure S2. Proportion of time spent vigilant for baboons in response to simulated predator auditory cues in Gorongosa National Park, Mozambique, across A) predator cues, B) year, C) habitat, and D) baboon age and sex class**. Panels B-D exclude data from playbacks of control (avian) cues. Observations in which the baboons fled from the cue immediately are excluded. Boxplots represent the median and interquartile range.

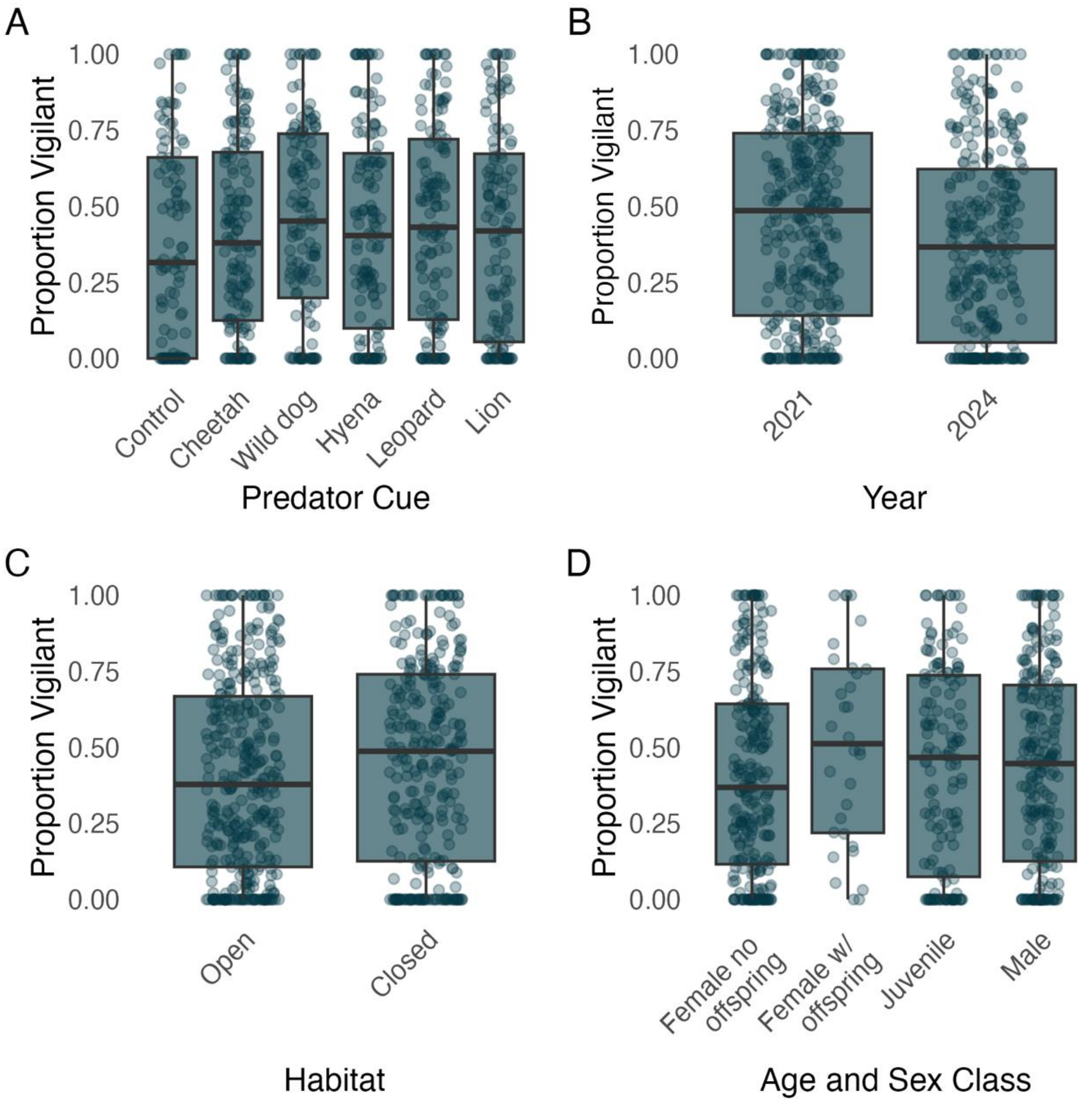

**Figure S3. Kaplan-Meier survival curves showing the probability of not fleeing throughout video duration in response to simulated predator audio cues in Gorongosa National Park.** Hatch marks indicate censored observations, representing trials in which no flight occurred before the observation period ended. Shaded areas indicate 95% CIs.

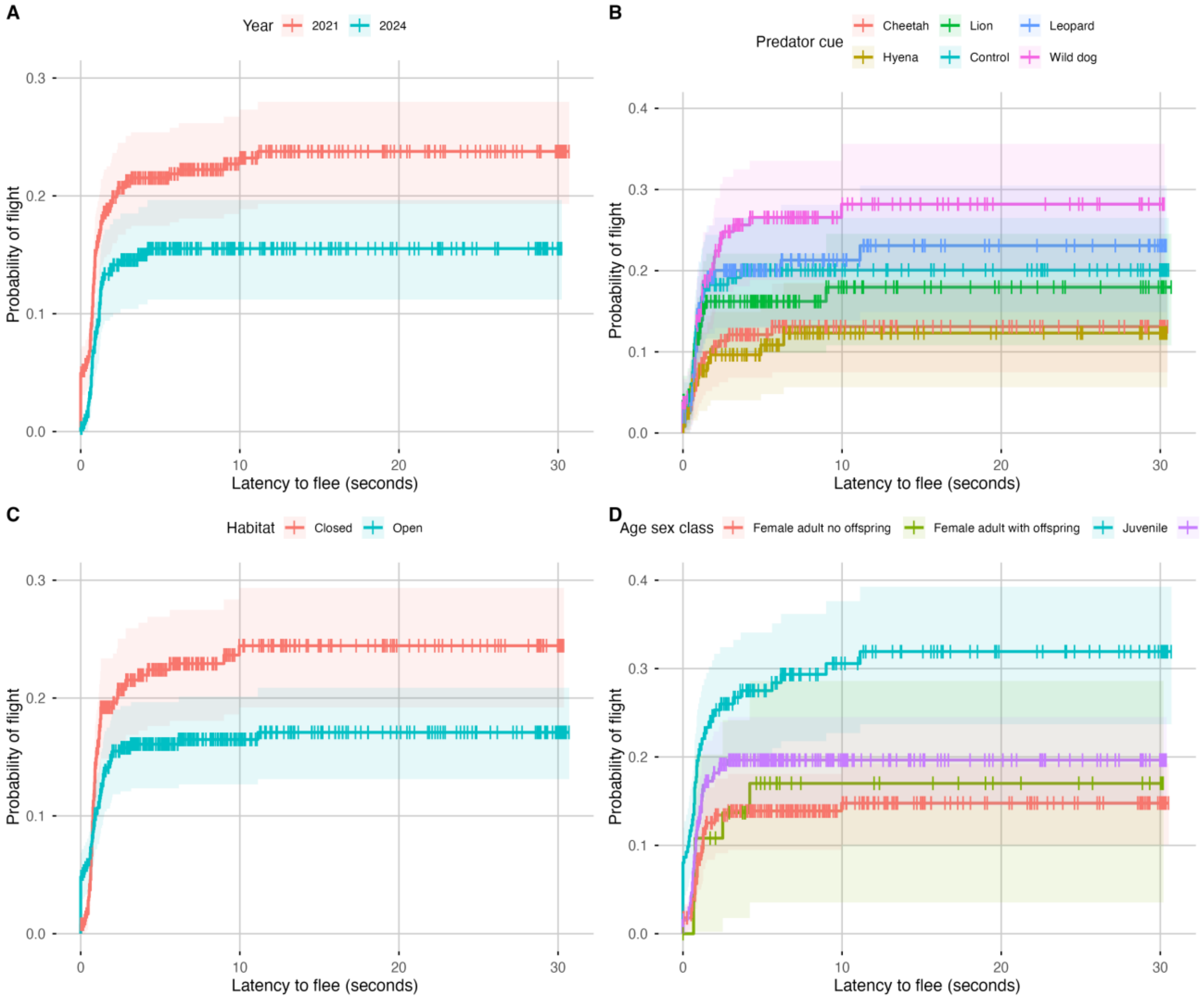

**Table S2. Ethogram for chacma baboon behaviors.** To be classified as exhibiting the behavior, an individual had to fully match the description.

| behavioral class | behavior | Description |
|---|---|---|
| Vigilant | Moving with vigilance | ● Exhibiting both vigilance (scanning or staring) simultaneously with moving with a four legged gait |
| Vigilant | Staring | ● Staring in one direction for greater than two seconds (Clarke et al., 2012)<br>● Accompanied by no movement in any direction<br>● Oriented towards ABR |
| Vigilant | Scanning | ● Visually scanning surrounding environment (Matsumoto-Oda et al., 2018)<br>● Head rotating by at least 45° (Clarke et al., 2012)<br>● May or may not be orienting towards the ABR |
| Vigilant | Standing and staring | ● Rising onto hind legs<br>● Staring in direction of source of sound |
| Vigilant | Startling | ● Fast, short acceleration away from source of sound<br>● Could be a short running burst or a jump |
| Non-vigilant | Moving non-vigilant | ● Slower pace of movement relative to fleeing<br>● All four limbs are involved in supporting body weight and movement forward (quadrupedal) (Druelle et al., 2021) |
| Non-vigilant | Feeding | ● Bringing hand to mouth and chewing (Clark et al., 2012)<br>● Peeling, digging, plucking, picking and holding (Segal, 2008) |
| Non-vigilant | Resting | ● Sitting, reclined, doing nothing (Clark et al., 2012) |
| Non-vigilant | Grooming | ● Licking oneself and using hands to clean (Clark et al., 2012) |
| Non-vigilant | Social interaction | ● Requires two or more baboons in close proximity at the start of behavior but can lead baboons separating. Requires initial proximity but doesn't have to end that way (ex. displacement by dominant) |
| Non-vigilant | Climbing | ● Quadrupedal climbing by supporting body weight with all four limbs while moving either horizontally or vertically across structures (such as trees or rocks) not on |

| | | |
|---|---|---|
| | | the ground (Preuschoft, 2002) |
| Flight | Fleeing | ● Faster, more dynamic movement away from source of sound compared to moving<br>● Usually a quadrupedal (four-legged) gallop or trot (Druelle et al., 2021)<br>● In this study, by the end of this behavior the individual will have left the field of view (frame) |
| Occluded | Occluded | ● An individual is hidden behind an obstruction (ex. Bushes, trees) or due to technical limitation and therefore their activity could not be discerned at a point in time (Kholiavchenko et al., 2024)<br>● Individuals head/face in no longer in frame |

**Table S3. Predictors of flight in baboons exposed to experimental predator cues in Gorongosa National Park, Mozambique**. Summary of model average regression coefficients and 95% CIs for the averaged model, and AIC summed model weights for each covariate. For the categorical variables, the reference categories are Year = 2021, Species = Cheetah, Habitat = Open, Class = Adult female (without offspring). ABR site was included as a random effect.

| Term | Model Average Regression Coefficient | 95% Confidence Interval | AIC summed model weights |
|---|---|---|---|
| Intercept | -2.053 | -2.933 – -1.174 | 1.00 |
| Age-sex class = Female adult with offspring | 0.304 | -0.690 – 1.299 | 0.999 |
| Age-sex class = Juvenile | 1.114 | 0.591 – 1.637 | 0.999 |
| Age-sex class = Male adult | 0.350 | -0.142 – 0.842 | 0.999 |
| Year = 2024 | -0.636 | -1.108 – -0.164 | 0.91 |
| Species = Hyena | 0.282 | -0.390 – 0.954 | 0.87 |
| Species = Leopard | 0.589 | -0.067 – 1.244 | 0.87 |
| Species = Lion | 0.709 | 0.057 – 1.360 | 0.87 |
| Species = Wild dog | 0.977 | 0.355 – 1.599 | 0.87 |
| Group size | -0.118 | -0.290 – 0.053 | 0.51 |
| Habitat = Closed | 0.415 | -0.153 – 0.983 | 0.46 |
| Day of study | -0.004 | -0.018 – 0.010 | 0.32 |

**Table S4. Predictors of proportion of time spent vigilant in baboons exposed to experimental predator cues in Gorongosa National Park, Mozambique**. Summary of model average regression coefficients and 95% CIs for the averaged model, and AIC summed model weights for each covariate. For the categorical variables, the reference categories are Year = 2021, Species = Cheetah, Habitat = Open, Class = Adult female (without offspring). ABR site was included as a random effect.

| Term | Model Average Regression Coefficient | 95% Confidence Interval | AIC summed model weights |
|---|---|---|---|
| Intercept | 0.398 | -0.022 – 0.819 | 1.000 |
| Day of study | -0.012 | -0.018 – -0.005 | 0.98 |
| Group size | -0.043 | -0.113 – 0.027 | 0.42 |
| Age-sex class = Female adult with offspring | 0.526 | 0.049 – 1.003 | 0.40 |
| Age-sex class = Juvenile | -0.030 | -0.299 – 0.238 | 0.40 |
| Age-sex class = Male adult | -0.016 | -0.248 – 0.217 | 0.40 |
| Year = 2024 | -0.151 | -0.437 – 0.135 | 0.39 |
| Habitat = Closed | 0.017 | -0.332 – 0.367 | 0.27 |
| Species = Hyena | 0.043 | -0.261 – 0.347 | 0.03 |
| Species = Leopard | 0.068 | -0.237 – 0.373 | 0.03 |
| Species = Lion | -0.073 | -0.382 – 0.237 | 0.03 |
| Species = Wild dog | 0.076 | -0.230 – 0.382 | 0.03 |

**Table S5. The top models for the probability of flight of baboons to simulated predator cues in Gorongosa National Park, Mozambique.** Only models with <2 delta AIC are included in this table, although all models were used in model averaging. The site ID was included as a random effect in each model.

| Model variables | df | logLik | AICc | delta AICc | weight |
|---|---|---|---|---|---|
| Age-sex class + Group size + Predator cue + Year | 11 | -334.94 | 692.23 | 0.00 | 0.16 |
| Age-sex class + Predator cue + Year | 10 | -336.10 | 692.51 | 0.27 | 0.14 |
| Age-sex class + Habitat + Predator cue + Year | 11 | -335.07 | 692.51 | 0.28 | 0.14 |
| Age-sex class + Group size + Habitat + Predator cue + Year | 12 | -334.15 | 692.72 | 0.49 | 0.13 |
| Age-sex class + Group size + Predator cue + Year + Day of study | 12 | -334.86 | 694.14 | 1.91 | 0.06 |

| **Model variables** | **df** | **logLik** | **AICc** | **delta AICc** | **weight** |
|---|---|---|---|---|---|
| Age-sex class + Group size + Predator cue + Year | 12 | -373.82 | 772.00 | 0.00 | 0.20 |
| Age-sex class + Group size + Habitat + Predator cue + Year | 13 | -372.89 | 772.22 | 0.21 | 0.18 |
| Age-sex class + Habitat + Predator cue + Year | 12 | -374.00 | 772.37 | 0.36 | 0.17 |
| Age-sex class + Predator cue + Year | 11 | -375.19 | 772.70 | 0.70 | 0.14 |

**Table S6. The top models for the proportion of time spent vigilant by baboons in response to experimental predator cues in Gorongosa National Park, Mozambique.** Only models with <2 delta AIC are included in this table, although all models were used in model averaging. The site ID was included as a random effect in each model.

| Model variables | df | logLik | AICc | delta AICc | weight |
|---|---|---|---|---|---|
| Day of study | 4 | 321.74 | -635.42 | 0.00 | 0.16 |
| Day of study + Group size | 5 | 322.30 | -634.51 | 0.90 | 0.10 |
| Day of study + Year | 5 | 322.23 | -634.36 | 1.05 | 0.10 |
| Day of study + Age-sex class | 7 | 324.18 | -634.19 | 1.23 | 0.09 |
| Day of study + Group size + Age-sex class | 8 | 325.20 | -634.17 | 1.24 | 0.09 |
| Day of study + Group size + Year | 6 | 322.84 | -633.54 | 1.88 | 0.06 |
